# Causal disconnectomics of motion perception: insights from TMS-induced BOLD responses

**DOI:** 10.1101/2022.03.03.482512

**Authors:** Estelle Raffin, Roberto F. Salamanca-Giron, Krystel R. Huxlin, Olivier Reynaud, Loan Mattera, Roberto Martuzzi, Friedhelm C. Hummel

## Abstract

Understanding how focal perturbations lead to large-scale network (re)organization is essential for accurately predicting the behavioral consequences of brain lesions. In this study, we applied a ‘virtual lesion’ approach by means of short bursts of 10 Hz transcranial magnetic stimulation (TMS) over either early visual areas (EVA) or the medio-temporal area (MT) in healthy participants, while acquiring concurrent functional MRI. TMS delivered during the early stages of motion processing selectively impaired direction discrimination at both sites, while global motion perception remained unaffected. These behavioral effects were accompanied by a common local increase in BOLD activity, but distinct patterns of network reorganization. Perturbation of EVA led to more robust and efficient functional adaptation, suggesting greater resilience to focal disruption. In contrast, behavioral impairments following MT stimulation were associated with a less organized, more random network structure. Together, these findings underscore the potential of TMS-fMRI coupling as a powerful approach for mapping *causal disconnectomics*—the dynamic relationships between localized neural disruption and widespread functional and behavioral outcomes providing a better understanding of lesion-induced brain changes in neurological disorders such as stroke.

**Highlights:**

- TMS-induced perturbation of the early visual areas (EVA) or the mediotemporal area (MT) area selectively impairs motion direction discrimination.
- The TMS perturbation is associated with a context-dependent local up-scaling of BOLD activity in both areas.
- The two visual areas display distinct topological networks’ adaptation in response to TMS, reflecting different levels of network resilience to a focal lesion.
- TMS-fMRI coupling can be used to assess ‘causal disconnectomics’ and to precisely map how a local perturbation propagates to large-scale behavioural deficits.

## 1. Introduction

Brain networks dynamically adjust their spatiotemporal organization to changing contexts and ongoing demands (Telesford *et al*., 2011; Papo *et al*., 2014). Such adaptive networks can be characterized by their topological transformation, their state-dependency and/or their state transition. The adaptability of brain networks is crucial for maintaining an optimal balance between information segregation and integration, ensuring functional stability, efficient local processing, and communication through long-range connections. Understanding how flexible and distributed brain networks give rise to complex behavior is a central challenge in systems neuroscience. This adaptability may enhance the resilience of functional networks in response to progressive neurodegeneration or focal lesions (Bassett & Bullmore, 2009; Bullmore & Sporns, 2012; Fornito *et al*., 2015). Today, a comprehensive understanding of the causal relationship between a focal lesion and brain network organization is still missing, leading to the inability to accurately predict resulting symptoms and recovery processes and develop targeted treatment strategies.

Directly testing the systematic association between a focal lesion and brain reorganization in humans is challenging, because of the difficulties in selectively altering activity *in vivo* and decipher the causal impact on brain networks. While advances in neuroimaging have provided detailed descriptions of functional and structural brain connectivity (Lim *et al*., 2019; Ma *et al*., 2022; Litwińczuk *et al*., 2022; Blanco *et al*., 2024), these correlational approaches offer limited insights into the causal mechanisms that underlie network dynamics and behavior. An emerging approach is to combine non-invasive brain stimulation (e.g., transcranial magnetic stimulation, TMS) with neuroimaging techniques (in particular functional magnetic resonance imaging (fMRI)) (Bergmann et al., 2021; Ozdemir et al., 2020; Vink et al., 2018, Raffin et al., in press). The combination of TMS with fMRI provides a unique opportunity to temporarily perturb a brain region and assess the changes in local brain activity and the resulting large-scale network responses in vivo that may underlie changes in task performance (Beckers & Hömberg, 1992; Ruff *et al*., 2006; Bestmann S., 2008; Ricci *et al*., 2012; Leitão *et al*., 2015; Cocchi *et al*., 2016). Several attempts have been made to combine fMRI with TMS to capture the impact that TMS can have on patterns of functional connectivity in specialized, large-scale, brain systems (Ruff *et al*., 2008; Jung *et al*., 2020). This holds great promise for establishing *individual causal connectomics* (Glick *et al*., 2024)— the ability to map the local effects of targeted perturbations of brain nodes and how they propagate across interconnected networks in individual participants.

Despite its conceptual appeal, key physiological mechanisms underlying TMS-fMRI responses remain incompletely understood. First, the local neural response to TMS reflects a complex interplay between intrinsic cortical excitability, synaptic connectivity, ongoing oscillatory activity, and network state. Second, the associated local TMS-evoked BOLD response and its physiological underpinnings remain to be fully elucidated (Rafiei & Rahnev, 2022). However, earlier studies have shown that these local perturbations can propagate through polysynaptic pathways, producing remote BOLD responses that seem to reflect both direct anatomical projections and state-dependent functional interactions (Ruff *et al*., 2006; Bestmann S., 2008; Siebner *et al*., 2022). The distinct propagation patterns across brain regions, combined with individual variability, offer a valuable window into mapping subject-specific causal connectivity—*causal connectomics*.

Describing personalized causal connectomics is particularly relevant when considering the application of TMS-fMRI to complex behavioral domains such as perceptual decision-making and motion direction discrimination. Motion direction discrimination is supported by a distributed network that integrates early visual areas (V1/V2) or EVA (Moore *et al*., 2001; Koivisto *et al*., 2010), motion-sensitive temporal regions (MT) (Newsome & Paré, 1988; Liu & Pack, 2017), the intraparietal sulcus involved in sensory evidence accumulation (Chen *et al*., 2017; Zhang *et al*., 2022; Wongtrakun *et al*., 2025), and prefrontal circuits supporting decision-making and response selection (Kennerley & Walton, 2011; Lin *et al*., 2020). Impairments in this system can arise from dysfunctions at multiple network levels, leading to behavioral deficits that may be difficult to attribute to any single region. Disentangling whether such deficits arise from primary sensory dysfunction, impaired integration, or altered top-down control requires tools that can interrogate how information flows across the network. By applying TMS-fMRI perturbations to key nodes within the motion discrimination network, here to the EVA and MT, it becomes possible to experimentally probe how focal disruptions alter distributed network dynamics and behavior. This approach may ultimately allow us to characterize individual-specific network vulnerabilities that give rise to multidomain symptoms, not only in healthy brain function but also in clinical populations where sensory, cognitive, and motor symptoms often co-occur (Siegel *et al*., 2016; Fleury *et al*., 2022; Jimenez-Marin *et al*., 2022). However, achieving such mechanistic understanding requires further efforts to relate region-specific TMS-evoked local and network-level responses to behavioral outcomes.

In this study, we aim to address these challenges by combining TMS-fMRI perturbation mapping targeting EVA and MT with behavioral assessments of motion direction discrimination, providing novel insights into the causal architecture of perceptual decision-making networks. These two regions were chosen because they allowed us to compare the whole brain consequences of a TMS perturbation affecting a primary integrative brain region, acting as a gateway to other ‘higher’ visual areas, and in contrast, a more functionally specialized brain region (Simoncelli & Heeger, 1998; Grossberg *et al*., 1999; Rust *et al*., 2006; Mineault *et al*., 2012). Simulation studies suggest that inhibitory rTMS targeting peripheral regions may exert more pronounced acute changes in network organization than stimulation of hub or integrative regions (Gollo *et al*., 2017). Accordingly, we hypothesized that, as a primary integrative hub, EVA would display greater local and network-level homeostasis following TMS, thereby attenuating the propagation of perturbation effects and limiting both behavioral impairments and widespread network reconfiguration compared to MT (Das *et al*., 2012; Gollo *et al*., 2017; Tu *et al*., 2021). Therefore, we anticipate that differences in TMS-induced behavioral impairments across behavioral state and across processing stages will be explained by distinct patterns of TMS signal propagation—shaped by the hierarchical network level of the targeted area—rather than by variations in local BOLD responses to TMS.

## 2. Methods

### Participants

In total, thirty healthy subjects (seventeen males: mean age 28.8 years, range 19–39 years) were recruited in the experiment. All participants provided informed written consent prior the experiment and none of them met the MRI or TMS exclusion criteria (Rossi *et al*., 2021). This study was approved by the local Swiss Ethics Committee (2017-01761) and performed in accordance with the Declaration of Helsinki.

### Experimental Design and Task Procedure

Sixteen participants (nine males, mean age 26.5 years, range 19-32 years) performed two TMS-fMRI sessions (except one drop-out for the second session). During the first session, TMS was applied to the right EVA (TMS_(EVA)_: mean(±SD) MNI coordinate: 8(5);-76(4);9(6)) using phosphenes perception when possible (in 9/16 participants) or the O2 position based on the 10-20 EEG system; during the second session, TMS was applied to the functionally defined right MT (TMS_(MT)_: mean MNI coordinates (SD): 46(4); −83(6); 11(5)) (Figure 1A). We chose this study design to maximize our chances to optimally target the individual MT area through a dedicated MT localizer acquired during the first session (see online supplementary materials for a description of the MT functional localizer). For the two online TMS-fMRI sessions, TMS was applied during a *motion direction discrimination (MDD) Task* and at *Rest*. These sessions had the same content except the anatomical T1-MPRAGE sequence and the MT functional localizer sequence, which were only performed during the first session. A short offline session prior the MT session was needed to locate the individual MT cluster using a neuro-navigation system (Localite GmbH, Bonn, Germany).

**Figure 1.**
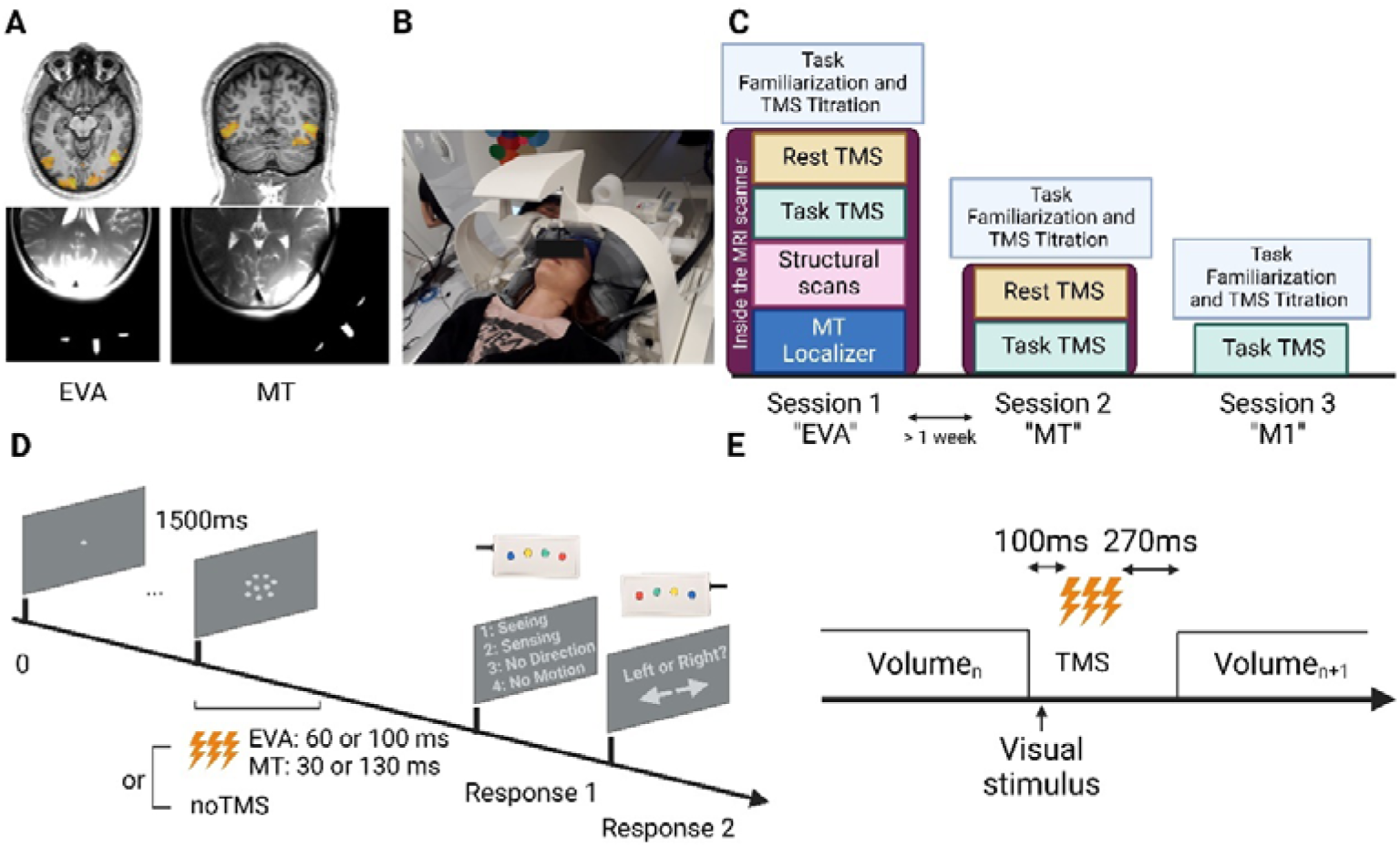
**A: TMS targeting.** TMS targets were determined by the functional localizer of EVA and MT for precise TMS positioning (see online materials for further details about the motion processing localizer). The two images on the right part show the T2 image of a representative subject with the oil capsules placed on the TMS-MRI coil casing to monitor the coil position located on the TMS coil for EVA and MT. These markers formed a “T-like shape” indicating the center of the coil as well as the coil orientation as seen on a T2 HASTE image acquired with the two TMS-compatible MRI coil arrays (56 axial slices, slice thickness = 0.8 mm, TR = 1500 ms, TE = 135 ms, FOV = 216 mm, flip angle = 140°). Coil positions were readjusted if needed. **B: MRI setup**. The MRI-TMS coil was positioned over the occipital area while the MR-receiving coil was positioned over the frontal lobe. Both were maintained via a vacuum pillow to ensure the absence of movement or vibration and to ensure a comfortable position for the participant. **C: Study timeline**: Description of the two experimental sessions; **D: One single trial of the motion discrimination task**. The visual stimuli involved a group of white dots in a 5° diameter circular aperture centered in the middle of the screen. The dots were either static or moving coherently to the right or to the left at a density of 2.6 dots per degree. Dots were displayed for 50, 67 or 84 ms (corresponding to 3, 4 or 5 frames). This duration was individually adjusted prior the fMRI measurement to ensure approximatively 70% accuracy in the noTMS trials. Just after the stimulus presentation, participants were asked to subjectively rate their perception using a response box on their right hand with a 1 to 4 scale, 1 being “I clearly saw the motion” (Seeing), 2 being “I sensed the motion” (Sensing), 3 being “I saw something moving but I’m unable to judge the direction” (No Direction) and 4 being “I didn’t see any motion” (No Motion) (Koivisto et al., 2010; Grasso et al., 2018). When the participants pressed the three first options, they were then asked to judge whether the dots moved to the right or to the left with a response box on their left hand. If participants pressed 4, the task moved to the next trial. E: Schematic of the BOLD sequence allowing artefact free combination of alpha TMS bursts.

An offline control experiment was performed to control for TMS non-specific effects on additional fifteen participants (seven males, mean age 28.9 years, range 22-39 years). We repeated the *MDD task* with TMS using the same setup (neuronavigation and eye-tracking systems) and the same task parameters (see below) but stimulating the right motor cortex (TMS_(M1)_, mean(SD) MNI coordinates: 53(7);-4(2);54(3), corresponding to the individual first dorsal interossei muscle’s motor hotspot of the participants), outside the MRI scanner. Figure 1B summarizes the study timeline.

For the three sessions, the *MDD task* consisted in rating subjective perceptual experience of visually presented motion stimuli and in performing a forced-choice direction discrimination task on the same moving stimuli (Koivisto *et al*., 2021). The visual stimuli involved a group of static or moving dots coherently to the right or to the left. During the familiarization phase outside the scanner, the duration of stimulus presentation was individually defined (either 50, 67 or 84 ms, corresponding to 3, 4 or 5 frames) to ensure approximatively 70% accuracy in the *noTMS* trials. The stimulus duration was then kept constant throughout the experiment (see Figure 1D for an illustration of the task).

Just after the stimulus presentation, participants were asked to rate their perception using a response box with their right hand using a 1 to 4 scale (see Figure 1’s caption for more details). In the main experimental conditions, TMS bursts were given to EVA when visual signals first reach the target region 60 ms after the onset of the visual stimulus (TMS_(EVA)_) (Lamme, 2001) and 30 ms after stimulus onset (TMS_(MT)_) for MT to account for the direct thalamic-extrastriate pathway (Beckers & Hömberg, 1992; Lamme & Roelfsema, 2000; d’Alfonso *et al*., 2002; Koivisto *et al*., 2010). This ensured similar perturbation of the feed-forward thalamic inputs to the two regions. Later TMS onsets were also tested in half of the trials, 100ms for EVA and 130ms for MT selectively interfering with late processing stages. The total duration of the TMS bursts was 200ms to cover the full window of visual processing in the two regions. TMS intensity was set to ≈ 80% [range: 75 to 90%] maximal stimulator output (MSO), corresponding to dI/dt of 119–169 A/ μs. The resulting E-field magnitude at hotspot (99.9-th percentile) corresponded to values between 73 and 112 V/m. This intensity was individually adjusted prior the measurement to ensure phosphene sub-threshold stimulation and progressively increased until participants report pain or discomfort. The intensity was then chosen to get reliable BOLD signal while preserving participant’s comfort. For the M1 control experiment, the same EVA timings were used and TMS was given sub-threshold to avoid motor evoked potentials elicitation.

Conditions randomly alternated between TMS versus noTMS trials, moving versus static trials throughout the task, 50 trials each, giving a total of 300 trials. An inter-trial interval (ITI) of approximatively 1.5 sec was used. The task was displayed in the scanner through a 44cm x 27cm LCD monitor at a 2.5m distance via a mirror mounted on the head coil, on a frame on top of the TMS-fMRI setup (Figure 1B). For the offline M1 control experiment, the task was displayed on a mid-grey background LCD projector (1024 x 768, 144 Hz). To exclude bad performance due to TMS-induced blinking and to make sure that participants were constantly looking at the fixation point, gaze and pupils’ movements were controlled in real time with an EyeLink 1000 Plus Eye Tracking System (SR Research Ltd., Canada) in the scanner and in the behavioral room, sampling at a frequency of 1000 Hz. Trials were aborted and redone if deviation exceeded 1° from the fixation point. The duration of the *MDD Task-TMS* sequence was on average 16.5 minutes (SD = 4.30 minutes) depending on individual reaction times.

A *Rest-TMS* sequence was used to study the effect of TMS intensity. An event-related design was used to map the effect of TMS bursts composed of three pulses at alpha (10Hz) frequency. Three conditions were pseudorandomized and counterbalanced across the run: high-intensity TMS (*HighTMS)*, low-intensity TMS *(LowTMS)* and no TMS *(noTMS),* with 25 repetitions of each condition with an inter-trial interval (ITI) of 6 seconds (covering 3 repetition times). *HighTMS* intensity corresponded to the intensity used during the MDD Task while *LowTMS* was set to ≈38 % [35 to 43%] MSO. Participants were asked to look at a fixation cross throughout the acquisition. The duration of the *Rest TMS* sequence was 9 minutes. Note that the results of the Rest-TMS sequence are presented in the supplementary materials.

### Behavioral data analysis

Considering static stimuli, the mean motion perception accuracy, reflecting how well participants could distinguish between moving and static stimuli, was extracted as the percent of correct answer relative to the total number of trials belonging to the same condition. Using moving stimuli, motion direction discrimination performances were extracted as the percent of correct Left-Right judgments relative to the total number of trials of that condition. For both scores, mixed ANOVA were built with factors Site (EVA, MT and M1) and TMS (Early onset, Late onset and NoTMS). Violation of sphericity was corrected using Greenhouse–Geisser corrections and Tukey’s multiple comparisons tests were applied for post hoc analyses when significant main effects or interactions were found.

### Magnetic Resonance Imaging & Transcranial Magnetic Stimulation

MRI images were acquired at the Magnetic Resonance Imaging facility of the Human Neuroscience Platform of the Fondation Campus Biotech Genève (FCBG, Biotech Campus, Geneva, Switzerland), using a Siemens Prisma 3 Tesla scanner (Siemens Healthineers, Germany).

### Standard MRI sequences acquisition

Anatomical images were acquired with a 64-channel head and neck coil using a 3D MPRAGE sequence (TR/TE = 2300/2.96 ms, flip angle 9°, 192 sagittal slices, matrix = 256, 1×1×1 mm^3^ resolution) covering the whole head. For retrospective distortion correction, static field mapping was performed with a double-echo spoiled gradient echo sequence (TR= 652 ms, TE1/TE2=4.92/7.38 ms, resolution 2.2×2.2×2.2 mm^3^, whole head coverage), generating two magnitude images and one image representing the phase difference between the two echoes.

### Combined TMS-fMRI sequences acquisition

For combined TMS-fMRI images, two dedicated coil arrays were used (Navarro de Lara *et al*., 2017). This setup consisted of an ultra-slim 7-channel receive-only coil array, which was placed between the subject’s head and the TMS coil (MRI-B91, MagVenture, Farum, Denmark) and connected to a MagPro XP stimulator (MagVenture, Farum, Denmark). A second, receive-only MR coil was positioned over Cz in the EEG 10-20 system to allow a full coverage of the participant’s brain.

The *Rest-TMS* and MDD *Task-TMS* sequences were acquired with a GE-EPI sequence using the same parameters: 40 axial slices, slice thickness = 2.2 mm, in-plane resolution = 2.2 mm, TR = 2000 ms, TE = 30 ms, FOV = 242 mm, flip angle = 67°, GRAPPA = 2, Multiband Factor (MB) = 2. A gap was introduced between consecutive EPI volumes in order to guaranty artefact free MR images after TMS stimulation (Navarro de Lara *et al*., 2015). For both rest and task TMS sequence, a single repetition time (TR=2000 ms) was therefore composed of 40 slices acquired during 1430 ms followed by a gap of 570 ms before the next volume acquisition. The synchronization of TMS pulse was carried out with an in-house script using Matlab (R2019).

Static field mapping was also performed with the TMS-MRI coils using the same double-echo spoiled gradient echo sequence (TR = 652 ms, TE = 4.92 and 7.38 ms, slice thickness: 2.2 mm, in-plane resolution = 2.2 mm, flip angle = 60°) that generates two magnitude images and one image representing the phase difference between the two echoes. Additional MRI sequences for coregistration of TMS-fMRI data are described in the supplementary materials.

### Preprocessing of fMRI images

Statistical Parametric Mapping software (SPM12, Wellcome Department of Imaging Neuroscience, UK) was used for data pre-processing and analysis of the four datasets (*Rest TMS* and *Task TMS* datasets, on both EVA and MT). Spatial distortions due to field inhomogeneities were removed from all EPI images using the Fieldmap toolbox from SPM12 (Andersson *et al*., 2001). Subsequently, all time-series were motion-corrected to the first volume using 6 degrees of freedom (3 rotations, 3 translations) and drifts in signal time courses were corrected as well. To improve the normalization procedure, two intermediate steps were added (see online materials for further details about the additional sequences used for coregistration purposes). A first co-registration was performed between the mean realigned and slice timing corrected image and the SSFP sequence acquired with the same MR coil (and thus the same spatial coverage). The resulting, co-registered image was once more co-registered to the SSFP sequence acquired with the MR coil integrated into the scanner (i.e., the body coil, thus preserving the contrast). The latter could then be easily co-registered to the high resolution T1 image acquired with the 64-channel head coil that covered the whole brain, and later transformed into standard MNI space using a segmentation-based normalization approach (Ashburner & Friston, 2005). Finally, the normalized images were spatially smoothed using a Gaussian kernel (4 mm, Full-width half-maximal). The realignment parameters estimated during spatial pre-processing for the *Rest TMS* and *Task TMS* dataset were introduced in the design matrix as regressors of no interest in order to prevent confounding activations related to minor head movements during scanning. The experimental event-related designs were convolved with the canonical hemodynamic response function, and the resulting models described in the following section were estimated using a high-pass filter at 128 s to remove low-frequency artefacts.

### fMRI Univariate Analysis

EVA and MT datasets were similarly analyzed using general linear models (GLM) to calculate individual contrasts. For the *Task TMS* sequence, a design matrix with TMS conditions (*TMS_early_*, *TMS_late_*, *noTMS*) and visual state (*Moving*, *Static*) was constructed using all trials. To disentangle the effects of a TMS onsets and visual stimuli, we built GLM with random effects at the second-level, by calculating a flexible factorial design, as implemented in SPM12, with the within-subject factors *TMS condition* (TMS_early_, TMS_late_) and *Visual stimulus* (Moving and Static).

To explore the local effects of TMS across conditions, mean beta values were extracted from the stimulated regions EVA and hMT+/V5 defined as a 8 mm radius sphere based on the results of the group-level analysis of the motion processing localizer (supplementary online materials) using the MarsBaR toolbox (Brett *et al*., 2002)(right EVA: 6, −83, 11 and right MT: 58, −66, −3). Repeated measure ANOVAs were built with TMS (*TMS_early_*, *TMS_late_*, *noTMS*) and visual stimulus (*Moving*, *Static*) as within-subject factors for each site of stimulation (EVA and MT).

For the *RestTMS* sequence, we defined a design matrix comprising three conditions (*HighTMS*, *LowTMS* and *noTMS*). A second-level GLM with random effects was designed using a flexible factorial design, with the within-subject factors *TMS intensity* (*TMS_High_, TMS_Low_* and *noTMS).* Beta values were extracted locally for each condition using the same ROI as previsouly described.

The statistical significance threshold was set to a height threshold of p < 0.001 uncorrected, at the voxel level and to that of p < 0.05 at the cluster level after false-discovery rate (FDR) correction.

### fMRI Multivariate Analysis

Independent Component Analysis (ICA) was used on the motion discrimination task data to estimate independent spatiotemporal functional networks from the data and to visualize networks specifically modulated by TMS. ICA uses fluctuations in BOLD signal to split it into maximally independent spatial maps or components, each explaining a unique part of variance from the 4D fMRI data. Then, all components have their specific time course related to a coherent neural signal potentially associated with intrinsic brain networks, artifacts, or both. The group ICA of fMRI Toolbox (GIFT version 3.0b; (Calhoun *et al*., 2001)) was used. First, it concatenates the individual data followed by the computation of subject-specific components and time course. Maximum Description Length (MDL) and Akaike’s criteria were applied to estimate the number of ICs in our data. Using principal component analysis, individual data was reduced. Then, the informax algorithm (Bell & Sejnowski, 1995) was applied for the group ICA and estimated 16 components. In order to improve the IC’s stability, the ICASSO was applied and run 20 times (Himberg *et al*., 2004). Over the 8 extracted network components, we focused on the TMS specific networks defined by the overlap between the activated clusters and the coordinates of the stimulation target. Based on these independent components, we then examined how each condition (Moving or Static) covaries with the TMS networks using the *temporal sorting* option in the GIFT toolbox. Temporal sorting consists in regressing the fMRI specific time course for each individual against the design matrix for the experimental conditions. The resulting betas weights represent the degree to which network was recruited by the conditions. For a network, positive and negative b weights indicate the level of network recruitment in each experimental condition. Paired-sample t-tests were used to identify networks that were significantly modulated by TMS at rest and during motion discrimination using p < 0.05.

### Brain Networks generation and Graph Theoretical Analysis

In addition to ICA, we used the graph theoretical network analysis toolbox GRETNA to image connectomics and construct the functional brain networks derived from the full Task TMS_(EVA)_ and TMS_(MT)_ dataset comprising all the conditions of interest (e.g., noTMS, TMS, motion or static stimuli) (Wang *et al*., 2015). The whole brain was divided into 90 cortical and subcortical regions according to Automated Anatomical Labeling (Tzourio-Mazoyer *et al*., 2002), and the mean time series for each of the 90 regions was extracted. Pearson’s correlation coefficients for each pair of regions were calculated for the mean time series of all of the 90 regions, and z-transformed using Fisher’s Z transformation. Subsequently, a positive binary undirected connection functional network was constructed according to a range of selected threshold of the relation matrix. As there is no defined standard for threshold selection in the construction of a binary connection brain functional network, sparsity was used as a range of correlation coefficient thresholds for correlation metrics. It was defined as the existing number of edges in a graph divided by the maximum possible number of edges. In accord with previous studies, we set both the sparsity and Pearson correlation threshold of the functional network to range from 0.05 to 0.5 (in 0.05 steps), resulting in a more efficient functional network than a random network with the number of artificial edges minimized (Achard & Bullmore, 2007; Zhao *et al*., 2017). Graph theoretical analysis was applied to assess the topological and organizational properties focusing on 3 graph theoretical metrics (Rubinov & Sporns, 2010). These metrics were: the clustering coefficient (γ), which measures how much neighbors of a node are connected to each other and is closely related to local efficiency; characteristic path length (λ), which is the average number of edges needed to get from any node in the network to any other node in the network and is inversely related to global efficiency; small worldness (δ), which reflects whether the network balances global integration and local segregation for efficient information processing (Latora & Marchiori, 2001). To compare graph theoretical network properties between EVA and MT, integration of the topological metric over all selected ranges of sparsity values was calculated using GRETNA. Paired t-tests corrected for FDR (p1.<1.0.05) were performed between the topological measures of EVA and MT networks.

## 3. Results

### 3.1 Behavioural results

We used chronometric TMS-fMRI coupling in healthy participants to perturb and map the effect of TMS over the right EVA (TMS_(EVA)_) and over the right MT (TMS_(MT)_) during a motion direction discrimination task. An additional offline experiment targeted the right primary motor cortex (TMS_(M1)_) in another subgroup of healthy participants.

Figure 2A shows the mean and individual motion perception accuracy (+/− SEM), reflecting how well participants could distinguish between moving and static stimuli when a TMS burst was applied over EVA (top row), MT (middle row) and over the control region M1 (bottom row). A mixed ANOVA revealed no main effect of TMS (F_(1,24)_ = 2.9, p = 0.084, ŋ^2^ = 0.02) and no significant TMS by Site interaction (F = 0.9, p = 0.452, ŋ^2^ = 0.031), suggesting that TMS has no consequence on motion perception in these three regions. However, there was a Site effect (F = 6.8, p = 0.004, ŋ^2^ = 0.16), likely driven by the overall better performances in the M1 session. The same model applied to the motion direction discrimination performances (Figure 2B) revealed a significant TMS effect (F = 22.012, p < 0.001, ŋ^2^ = 0.2) and Site effect (F = 20.57, p < 0.001, ŋ^2^ = 0.2). There was also a TMS by Site interaction (F = 9.87, p < 0.001, ŋ^2^ = 0.124). When TMS was applied to EVA, performances were markedly affected by TMS_early_ compared to noTMS (t_(15)_ = 5.2, adjusted p < 0.001) and compared to TMS_late_ (t_(15)_ = 4.4, adjusted p = 0.034). Although slightly lower, the difference between noTMS and TMS_late_ was not significant (t_(15)_ = 1.45, p = 0.35).

**Figure 2:**
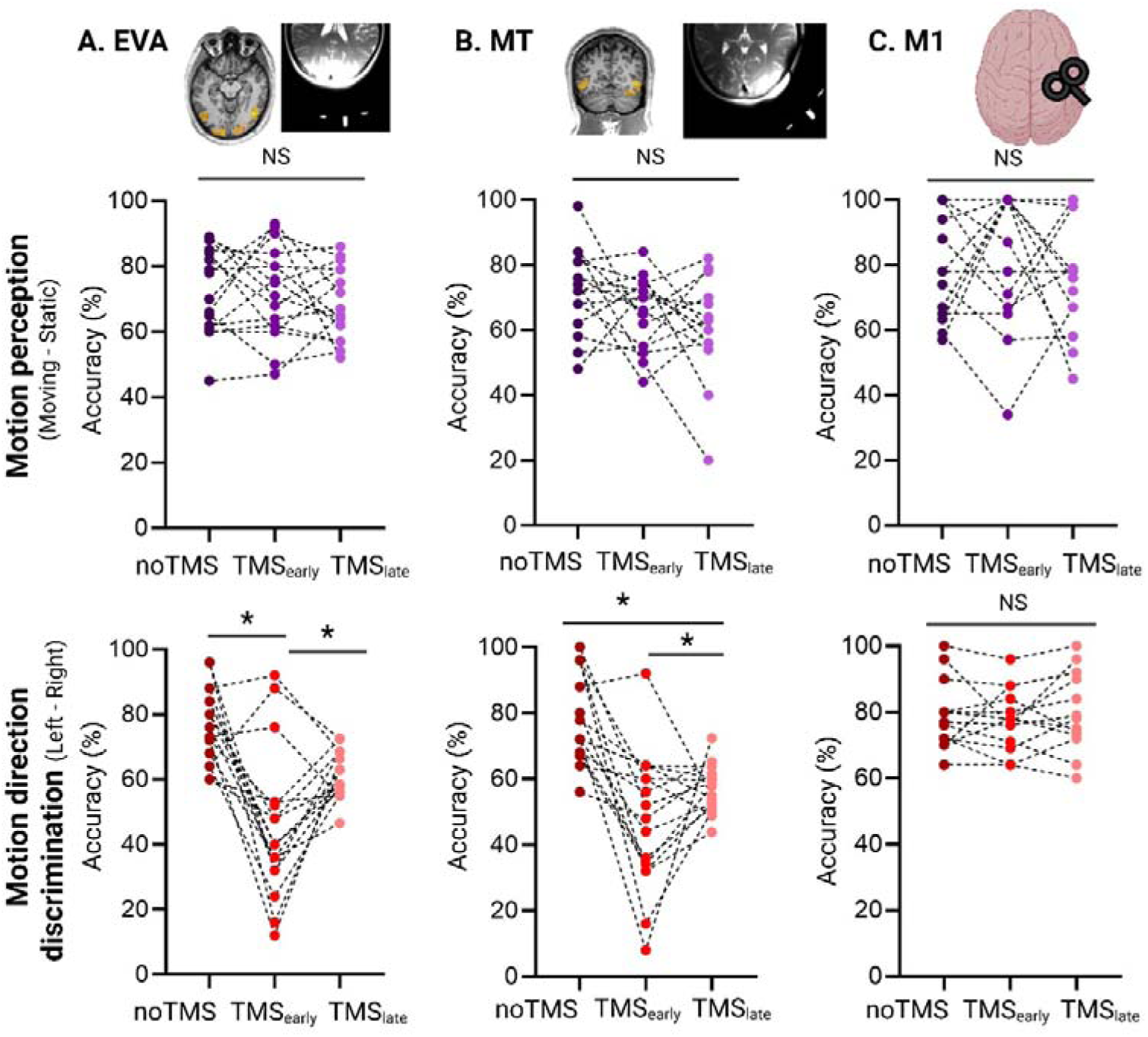
The upper row illustrates the individual data of motion perception under noTMS, TMS_early_ and TMS_late_ for TMS_(EVA)_ (A), TMS_(MT)_ (B) and TMS_(M1)_ (C); The bottom row illustrates the individual data of motion direction under noTMS, TMS_early_ and TMS_late_ for TMS_(EVA)_ (A), TMS_(MT)_ (B) and TMS_(M1)_ (C);

In contrast, when applied to MT, TMS_early_ and TMS_late_ negatively impacted performance compared to noTMS (TMS_early_: t_(15)_ = 5.2, adjusted p < 0.001, TMS_late_: t_(15)_ = 5.2, p < 0.001). After correction for multiple comparisons, performance under TMS_early_ and TMS_late_ were not different (t_(15)_ = 2.2, p = 0.1). Performances were similar in the three TMS conditions for M1 (all comparisons p>0.05) and baseline performances (noTMS conditions) were not different across sites (all comparisons p>0.05).

We also examined the relationship between awareness and accuracy by taking into account motion awareness ratings (see supplementary Figure S1). Without TMS, motion direction discrimination increased (moved away from chance level) as subjective awareness of motion increased. This indicates that without TMS, performance is tied to conscious motion awareness. The rmANOVA with TMS and Rating as within subject factors only revealed a significant Site by TMS condition interaction, confirming the previous analysis (F = 15.1, p < 0.011, ŋ^2^ = 0.09). There was no main effect of Awareness nor interactions with any other factor (all p>0.05).

In summary, TMS bursts exerted negative effects on motion direction discrimination at the early and late onsets in MT. TMS bursts were only efficient in disturbing motion directions discrimination in the early onset for EVA. The two onsets did not affect performances when targeting the control region M1.

### 3.2 Whole-brain and local responses to TMS perturbation

Next, we analyzed local BOLD activity in response to TMS bursts over EVA and MT applied during behavioral perturbation of motion direction discrimination. First, Figure 3A (TMS_(EVA)_) and Figure 3B (TMS_(MT)_) report the results of the average effects of TMS conditions. TMS_(EVA)_ induced significant clusters in the bilateral occipital cortex, the right superior and middle temporal gyrus, in the bilateral anterior and middle cingulate cortex, the left middle frontal gyrus, the bilateral caudate nucleus and in the right posterior thalamus. When TMS was given over the right MT, the average effect of TMS conditions returned significant clusters in the right occipital cortex, the right inferior and frontal gyrus, the right intra-parietal sulcus and the right cerebellum (cruz VI). These clusters overlapped with the TMS-induced activation at rest (see supplementary results and Table S1).

**Figure 3.**
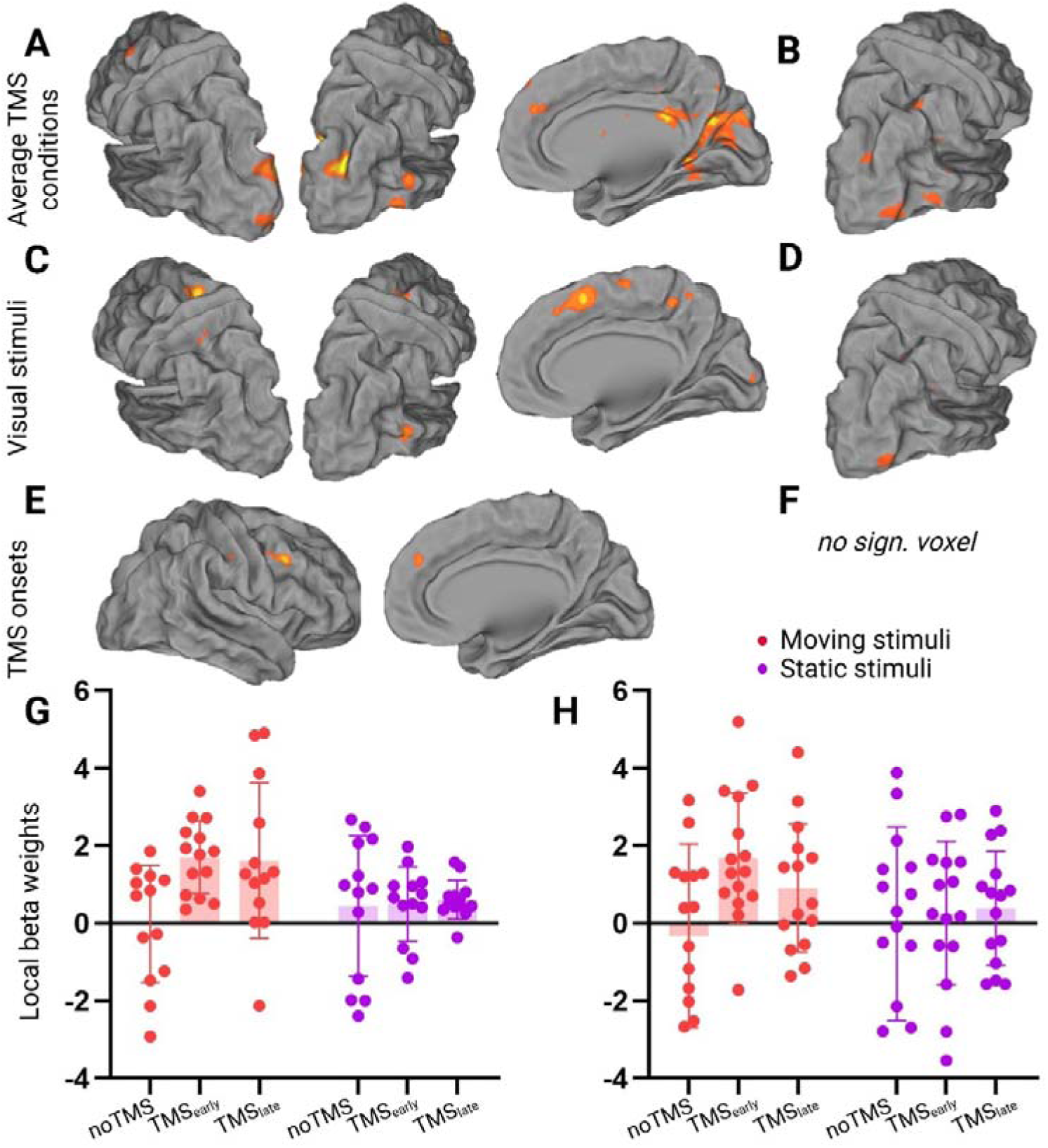
A: Activation maps corresponding to the average TMS conditions for TMS_(EVA)_; **B:** Activation maps corresponding to the average TMS conditions for TMS_(MT)_; **C:** Main effect of visual stimuli for TMS_(EVA)_; **D:** Main effect if visual stimuli for TMS_(MT)_; **E:** Main effect of TMS onsets for TMS_(EVA)_; **F:** Main effect of TMS onsets for TMS_(MT)_; **G:** Local beta weights extracted for the EVA ROI: **H:** Local beta weights extracted for the MT ROI. P(FWE)) < 0.001 at the cluster level.

We then examined the main effect of visual stimuli (difference between TMS applied during a moving or static stimuli) returned significant clusters in the right middle temporal gyrus and in the bilateral middle frontal gyrus the right superior frontal gyrus, the posterior cingulate cortex, and the right occipital cortex when EVA was stimulated (Figure 3C) and only in the right middle temporal gyrus, when MT was stimulated (Figure 3D). When stimulating the EVA, the main effect of TMS onsets revealed a significant cluster in the right middle frontal gyrus and in the right superior frontal gyrus (Figure 3E). There was no significant cluster when stimulating MT (see Supplementary Table S2 and S3 for the MNI coordinates of the significant clusters of all effects).

The extracted beta values from the stimulated EVA and MT regions were entered into an ANOVA with the factors TMS (noTMS, TMS_early_ and TMS_late_) and Visual stimuli (moving, static) and stimulation site (EVA, MT). This analysis revealed a significant TMS effect (F_(2,22)_ = 3.8, p = 0.038), mostly driven by an increase in beta weights for the two TMS conditions compared to no TMS, while there was no difference between the early and late TMS onsets. Finally, there was a significant TMS by Visual stimuli interaction (F_(2,22)_ = 6.1, p = 0.008) showing that the increase in activity only occurred in the combination early TMS associated with moving stimuli (noTMS, Moving vs early TMS, Moving: t_(12)_ = −4.2, p = 0.003 and early TMS, Static vs early TMS, Moving: t_(12)_ = −3.6, p = 0.015, supplementary Table S4 for all posthoc comparisons). There was no stimulation site effect, demonstrating that the local response to TMS were similar in both regions.

### 3.3 Network changes in response to TMS perturbation

We further investigated whether and how long-range brain activity was modulated by TMS during motion direction discrimination. We first applied an independent component analysis to extract hidden spatiotemporal components contained within the TMS_(EVA)_ and TMS_(MT)_ task data, returning a measure of functional connectivity. We then applied temporal regression analysis on the components’ time courses to compare network activity associated with the different experimental conditions.

The ICA applied to the TMS_(EVA)_ data during motion discrimination revealed 8 independent networks, among which 7 were considered as functional brain networks. Five networks (IC1, IC3, IC4, IC7 and IC8) were labelled as TMS-related networks because they overlapped with the stimulation site (Figure 4A and Supplementary Table S5 for the MNI coordinates of the IC). IC1, IC7 and IC8 covered the bilateral EVA while IC3 only encompassed the right EVA and IC4, the left EVA. Only IC1 and IC7 were significantly modulated by TMS as revealed by one-way ANOVAs modeling the t scores obtained from the temporal regression analysis on the components’ time courses for moving and static stimuli. IC1 was significantly down regulated by both TMS onsets when associated with moving stimuli only (one-way ANOVA: F_(2,15)_ = 3.6, p = 0.04, post hoc TMS_early_ vs noTMS: t_(15)_ = 1.9, p = 0.07, TMS_late_ vs noTMS: t_(15)_ = 2.2, p = 0.04). IC7 showed a different pattern of modulation reflecting a significant over-expression of the bilateral EVA only during TMS_early_, for moving (F_(2,15)_ = 3.5, p = 0.045, post hoc TMS_early_ vs noTMS: t_(15)_ = 2.4, p = 0.03) and static stimuli (F_(2,15)_ = 4.4, p = 0.03, post hoc TMS_early_ vs noTMS: t_(15)_ = 2.3, p = 0.035). All the other ANOVAs were not significant.

**Figure 4.**
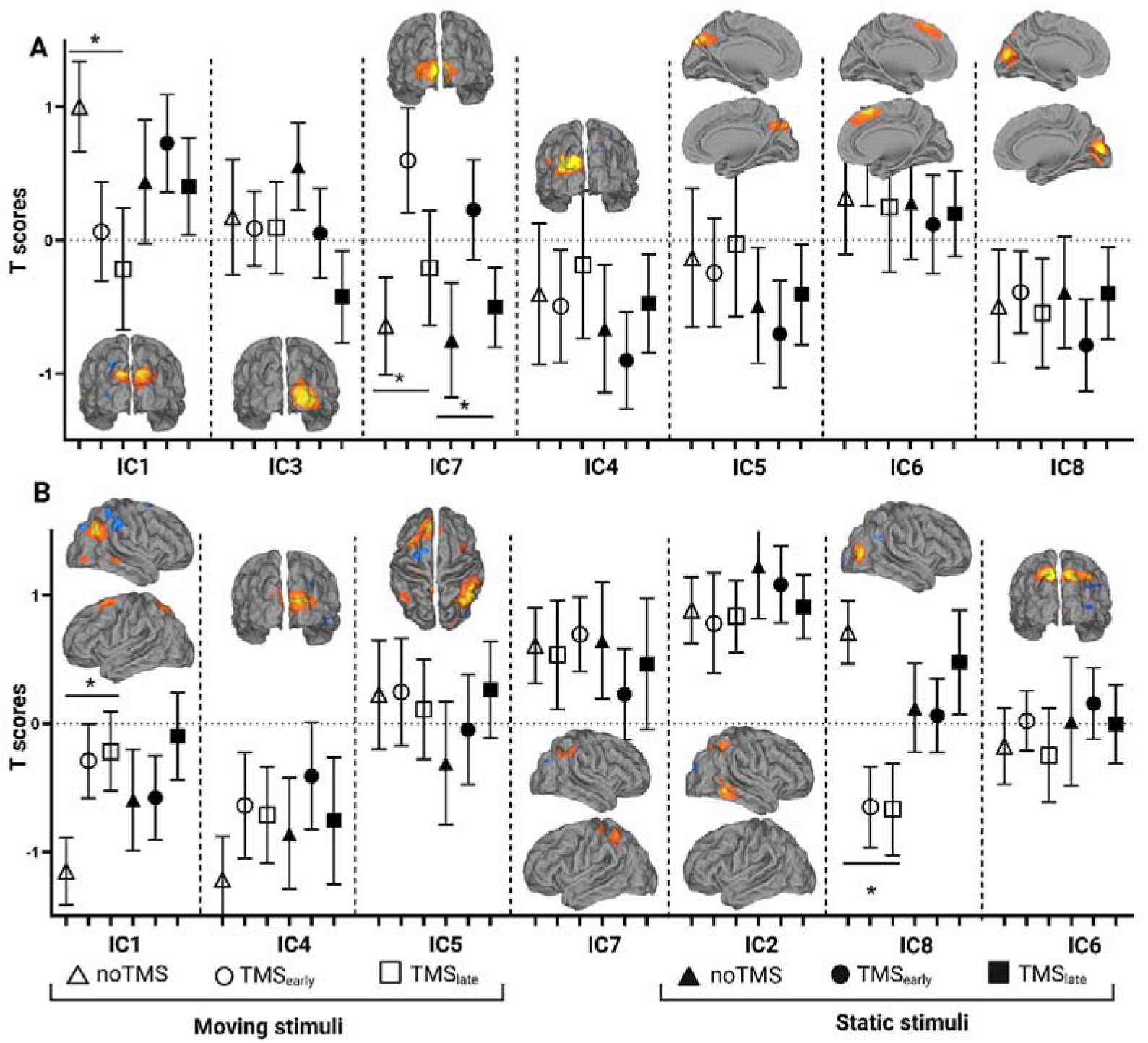
**A:** TMS_(EVA)_-specific component networks that are modulated by TMS during motion discrimination and temporal regression analysis for the noTMS and TMS_(EVA)_ conditions associated with moving or static stimuli; **B:** TMS_(MT)-_specific component networks that are modulated by TMS during motion discrimination and temporal regression analysis for the noTMS and TMS_(EVA)_ conditions associated with moving or static stimuli; Note that the components are sorted based on the percent of explained variance.

When TMS was given over MT, 7 components were considered as functional brain networks. Among them, three networks (IC1, IC2 and IC8) were labelled as TMS-related networks (see Supplementary Table S6 for the MNI coordinates of the IC). Interestingly, these networks were not only restricted to the right MT. IC2 and 8 also included the IPS and superior parietal cortex and the medial frontal gyrus. IC 1 activity was significantly enhanced by both TMS onsets when stimuli were moving (F_(2,15)_ = 3.6, p < 0.001 post hoc TMS_early_ vs noTMS: t_(15)_ = 2.7, p = 0.02, TMS_late_ vs noTMS: t_(15)_ = 3.8, p = 0.002) while IC8 was significantly depressed during moving stimuli (F_(2,15)_ = 7.4, p = 0.003, post hoc TMS_early_ vs noTMS: t_(15)_ = 3.2, p = 0.007, TMS_late_ vs noTMS: t_(15)_ = 3.1, p = 0.008, Figure 4B). All the other ANOVAs were not significant.

Finally, we used a graph theoretical analysis to assess the topological and organizational properties of these networks, while being agnostic to the TMS condition and visual stimulus. We first investigated small-world topological properties especially because networks with high small-world properties are believed to play a role in protecting the brain from local perturbation or damage. Both EVA and MT networks exhibited small-world properties (λ1.≈1.1 and σ1.>1.1) among the selected sparsity values (see Methods section). Interestingly, there was a significant difference in the small-world network parameters (Sigma) between EVA networks and MT networks with higher Sigma values for EVA networks at the sparsity values 0.3, 0.4, 0.45 and marginally at 0.35 (Figure 5A, left panel and supplementary Table S7 for all paired t-tests), also considering the mean across all sparsity values (t_(12)_ = 2.4, p = 0.034) (Figure 5B, left panel). There was no difference in normalized clustering coefficient (Gamma) (t_(14)_ = 0.42, p = 0.68) (Figure 5A & B, middle panels and supplementary Table S8 for all paired t--tests). Finally, characteristic path length (Lambda) was significantly higher at sparsity thresholds of 0.2, 0.25 and marginally at 0.15 for MT networks (Figure 5A, right panel and supplementary Table S9 for all paired t-tests). The difference between EVA and MT+/ networks was also significant when averaging across scarcities (t_(14)_ = 2.4, p = 0.03, Cohen’s d: 0.75) (Figure 5B, right panel).

**Figure 5.**
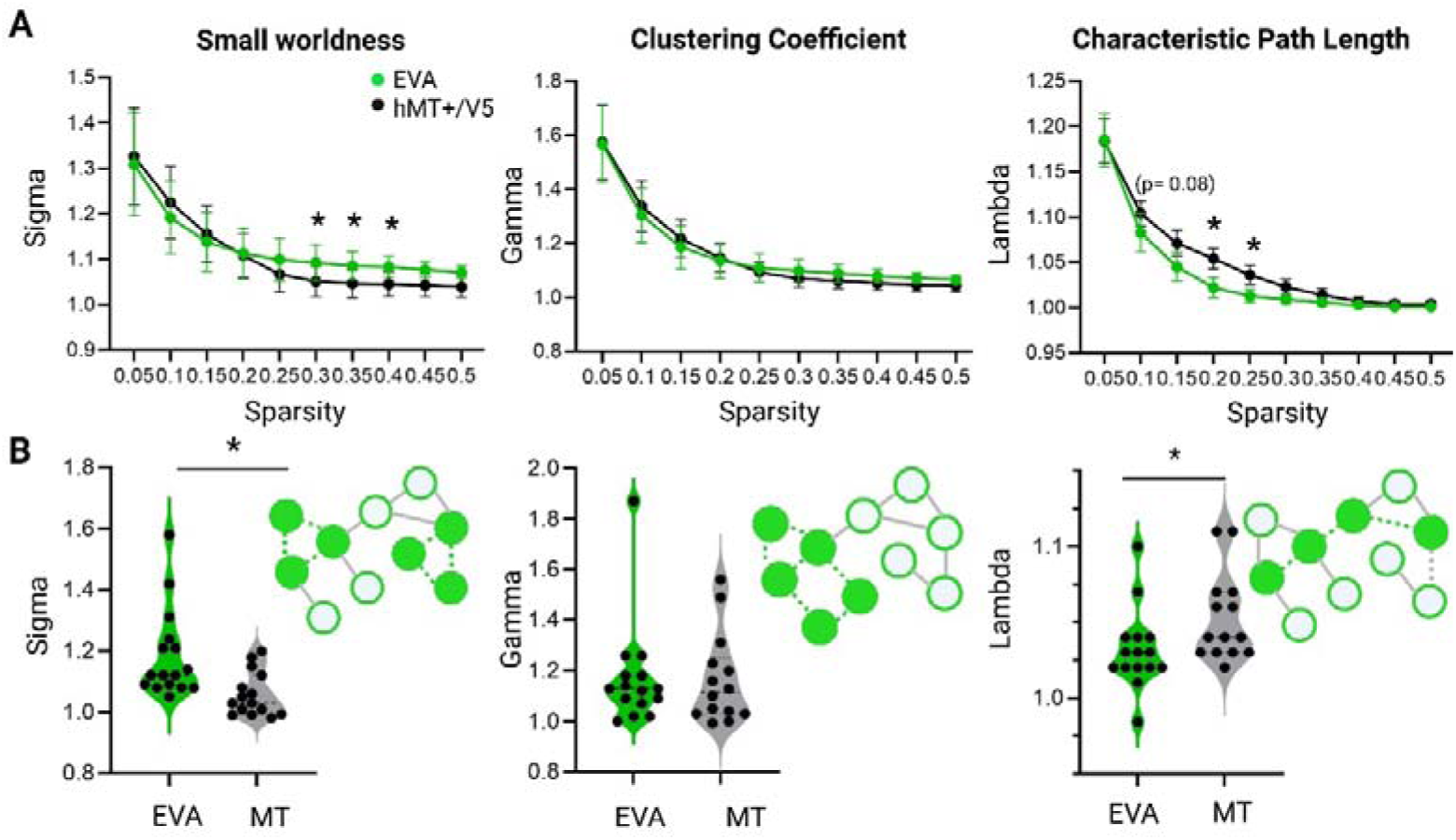
**A:** Small-worldness properties (left panel), normalized clustering coefficients (middle panel) and normalized characteristic path length (right panel) for TMS_(EVA)_ and TMS_(MT)_ along the sparsity thresholds of 0.05-0.50; B: Associated mean values of small-worldness properties (left panel), normalized clustering coefficients (middle panel) and normalized characteristic path length (right panel) for TMS_(EVA)_ and TMS_(MT)_ * p < 0.05.

### 3.4 Predicting TMS effects on behaviour with BOLD activity

To explain the effect of TMS on motion direction discrimination, we performed a series of multiple linear regressions using for each of the four conditions (TMS_(EVA)_early, TMS_(EVA)_late, TMS_(MT)_early, TMS_(MT)_late), the associated local beta scores, the t scores derived from the temporal regression of the significant TMS-ICs and the three graph theory metrics as fixed effects. The significant model explained the drop in performance associated with TMS_(MT)_early with Gamma values from the graph theory analysis as confirmed by the ANOVA on the model’s parameters (F_(1,6)_ = 6.6, p = 0.04), suggesting that TMS exerted a negative impact especially in participants showing lower cluster coefficient, implying a more random or decentralized network structure. All other models were not significant (see supplementary Table S10 for all the multiple linear regressions results).

## 4. Discussion

The goal of this study was to examine whether TMS perturbation elicits inter-individual differences in BOLD responses, potentially reflecting the magnitude of the induced behavioral impairment. We used short bursts of 10Hz TMS to disrupt either a low-level visual area (EVA) or a highly specialized, motion sensitive, visual area (MT) during concurrent fMRI. We first confirmed the causal involvement of both regions in motion direction discrimination, with more pronounced behavioral impairments following stimulation of MT. Our main findings were: (1) a local, context-dependent increase in BOLD activity common to both targeted areas; (2) distinct patterns of network reconfiguration; and (3) a relationship between TMS-induced performance decline after MT stimulation and a shift toward a more random network structure. These results suggest that TMS perturbation triggers area-specific topological adaptations, which may reflect differing levels of network resilience to focal disruption.

### 4.1. TMS-induced behavioural deterioration

In line with the concept of “virtual lesion” described in the literature (Pascual-Leone *et al*., 1999; Beynel *et al*., 2019), TMS bursts applied to either EVA and MT, resulted in a selective and transient deterioration of performance of a left/right motion direction discrimination task (except for the late TMS_(EVA)_ condition). This was interpreted to reflect successful causal and selective disruption of brain functioning. Interestingly, participants remained able to dissociate moving from static stimuli. This dissociation is likely supported by the existence of independent substrates mediating unspecific motion processing versus global motion direction integration in EVA and MT (Simoncelli & Heeger, 1998). There is evidence showing that a sub-population of neurons in MT focuses on the processing of local motion signals similar to that in EVA, and some other neuronal populations rather process global motion (Born & Tootell, 1992). Our TMS perturbation might have specifically impacted on the neurons in charge of motion directions decoding. The absence of significant impairment induced by the late onsets of TMS_(EVA)_ might be explained by different time delay of feedback loops in the motion discrimination system (Foss & Milton, 2000; Joukes *et al*., 2014). Considering now the relationship between motion awareness and motion direction discrimination (please see supplementary Figure SX), we found that the direction of the moving trials that were judged as “moving”, were not correctly guessed when both areas were stimulated. In contrast, “unaware” trials were unchanged. We can speculate that this residual, unconscious ability of detecting changes in the dots’ positions might be at least partially supported by the activation of the fast, direct geniculo-extra-striate pathway (Bridge *et al*., 2008; Ajina *et al*., 2015; Abed Rabbo *et al*., 2015).

### 4.2. Local TMS-induced activity

Our results demonstrate that short bursts of TMS can elicit local BOLD activation beneath the coil in both EVA and MT, even at relatively low intensities—below the phosphene threshold. While the presence of local BOLD activity elicitation in response to TMS is still unresolved in the field (Rafiei & Rahnev, 2022), we specifically found a local over-activation in the presence of moving visual stimuli for both regions. This coupling between local BOLD increase and decreased performance suggests that the additional activity may reflect the accumulation of task-irrelevant neural noise (Ruff *et al*., 2009; Bancroft *et al*., 2014). Such a mechanism aligns with established models of TMS-induced disruption, where stimulation is thought to activate neuronal populations that are not optimally tuned to the task-relevant feature—in this case, motion direction (Silvanto & Pascual-Leone, 2008). As a result, the effective signal-to-noise ratio within the motion-sensitive network would be reduced, impairing accurate motion discrimination. It also highlights the state-dependent nature of TMS effects: stimulation during active motion direction processing leads to qualitatively different outcomes than during static stimuli presentation, suggesting that the excitability and engagement of the targeted network at the time of stimulation critically shape the neuronal consequences. These findings reinforce the idea that TMS does not simply “excite” or “inhibit” local tissue, but interacts dynamically with ongoing neural activity (Romero *et al*., 2019; Siebner *et al*., 2022). This, together with other recent findings (Perera *et al*., 2024; Luo *et al*., 2025), might open the door to more tailored or context-sensitive brain stimulation protocols.

### 3. TMS induced whole-brain activity

An important question is whether this “local noise injection” propagates in the brain to destabilize larger functional networks. TMS-induced changes in BOLD activity were observed not only locally in the stimulated regions, but also in remote areas through functional coupling. For both EVA and MT stimulation, the spread of activity to distal regions—such as the intraparietal sulcus (IPS), frontal eye fields (FEF), and medial prefrontal cortex—replicates previous findings from TMS-fMRI studies (Ruff *et al*., 2006; Caparelli *et al*., 2010; Leitão *et al*., 2015). Notably, many of these remote regions are task-relevant and are known to be engaged during motion perception and direction discrimination (Offen *et al*., 2010; Sani *et al*., 2021), suggesting that the ongoing task context may also shape the propagation of TMS-evoked activity through the actively engaged network.

When TMS was applied during motion processing, additional BOLD activation was observed in regions such as the medio-temporal cortex (including MT) and medial frontal areas. Importantly, this increased activation coincided with a measurable decline in behavioural performance on the motion direction discrimination task, as well as with participants’ subjective reports of increased task difficulty. These findings suggest that TMS may disrupt efficient network integration during perceptual processing, potentially by altering the balance of activity within task-relevant networks. The differential effect induced by the early and late TMS onsets was only visible when stimulating EVA and manifested as an increase in activity in the right FEF and in the medial prefrontal cortex with the late onsets. These regions are involved in perceptual decision making and this time could reflect a compensatory mechanism that maintain stable performance (Imani *et al*., 2021).

### 4. TMS-induced network activity

To provide both a qualitative and quantitative framework for comparing network activity across different TMS-induced perturbation, we combined group Independent Component Analysis (ICA) with graph-theoretical metrics. ICA identified multiple functional networks, some directly linked to TMS and others more domain-general. Across both targeted regions (EVA and MT), the decomposition revealed components with divergent responses to TMS. In the case of EVA, for example, IC7— spatially overlapping with the site of stimulation—showed increased activation specifically following early TMS onset. In contrast, IC1, located more rostrally, exhibited a pattern of downregulation under the same stimulation condition. The opposing responses may reflect a balance between excitatory and inhibitory processes, possibly mediated by intra-areal or feedback projections (Shao & Burkhalter, 1996; Schwabe *et al*., 2006). Furthermore, the specificity of IC7’s response to early TMS onset raises questions about the temporal dynamics of network susceptibility, with different networks or nodes having critical windows of “vulnerability” or influence (Huang & Yu, 2017; Kottaram *et al*., 2018). This dissociation suggests that ICA successfully captured the complex, spatially and temporally differentiated dynamics of TMS-induced responses. It highlights ICA’s sensitivity not only to spatial localization but also to the functional state of the network—whether it is task-relevant, engaged, or modulated by TMS in a context-dependent manner. It further emphasizes that TMS effects are not uniform within a target region but depend on the local functional architecture and current cognitive or perceptual state.

Regarding network activity induced by TMS_(MT)_, we found that TMS engaged more spatially distributed networks, involving additional associative regions such as the FEF, the posterior parietal cortex (PPC), and medial prefrontal cortex (mPFC). In contrast, all but one of the independent components associated with TMS_(EVA)_ were largely confined to the stimulated visual area, indicating more localized network engagement.

To quantitatively assess these differences, we applied graph-theoretical analyses and observed distinct whole-brain network configurations depending on the stimulation site. Specifically, networks following TMS over EVA exhibited higher small-worldness—a topological property characterized by strong local clustering and short path lengths— compared to those following TMS over MT. This network architecture is thought to support efficient parallel information processing by maximizing information transfer at minimal wiring cost (Achard & Bullmore, 2007). The clustered, locally efficient organization of the TMS_(EVA)_ network may promote functional resilience and flexible redistribution of activity. This could help explain the relative preservation of behavioral performance during late TMS onset, suggesting that the network maintains coherence despite transient perturbations (Wu *et al*., 2020). In contrast, the more dispersed TMS_(MT)_ network configuration likely reflects broader propagation of task-specific interference across higher-order visual regions, potentially leading to a less efficient, more vulnerable network state. Congruently, the variable that best accounted for the TMS-induced performance impairments following TMS_(MT)_ was the clustering coefficient. This measure reflects the tendency of nodes within a network to form tightly interconnected clusters. High clustering values indicate a well-organized, locally efficient network structure, whereas lower values reflect a more random or fragmented configuration, consistent with patterns observed in our ICA analyses. Our findings showed that participants who exhibited greater performance declines had lower clustering coefficients, suggesting that a more dispersed network organization is detrimental to motion direction discrimination. It would be valuable in future work to investigate whether these network disruptions extend to other visual subfunctions—or even domain-general cognitive functions—in order to determine the broader functional impact of focal perturbations on brain-wide network dynamics.

This result fits with our initial hypothesis which was that EVA, given its role as an early-stage integrative cortical area, would maintain more stable synaptic and network dynamics in response to TMS, resulting in greater resistance to externally induced disruptions in both perceptual performance and functional connectivity compared to MT. It is in line with a modelling study, showing that brain hubs, where activity is integrated and further distributed (like EVA in our case) operate in a slower regime and appear to be functionally less affected by focal perturbations (Gollo *et al*., 2017). In contrast, perturbations of peripheral or more associative areas (e.g., MT), might have greater impact on acute network activity. Lesion studies also provided evidence that damage to the homologue of MT in behaving cats impairs more relearning of motion discrimination and learning transfer than lesions to EVA (Das *et al*., 2012). To sum up, the network analyses revealed differences between the two disrupted areas, not observable with classical activation analyses. It highlights the relevance of graph-theoretical metrics in capturing functionally meaningful changes in network topology. In particular, the clustering coefficient appears to be a sensitive marker of TMS-induced network reconfiguration and its behavioural consequences.

### 5. Limitations of the current work and open questions

The aim of this study was to develop a causal connectomic framework for understanding motion direction discrimination by selectively perturbing two key network nodes using TMS. However, it is important to recognize certain methodological limitations inherent to this approach. Unlike brain lesions, TMS-induced perturbations are both temporally transient and spatially focal. More critically, the physiological consequences of a lesion might differ fundamentally from those of TMS. Brain lesions not only disrupt local neuronal activity but also provoke widespread, often nonspecific responses—particularly in the hyperacute phase—such as excitotoxicity, inflammation, and diaschisis (Lai *et al*., 2014; Farooqui *et al*., 2022). These processes can profoundly alter interregional communication through mechanisms unrelated to normal synaptic signaling.

In contrast, the type of TMS used in this study induces neuronal firing and inter-areal communication, though likely mediated by different—and possibly more superficial or excitatory—neuronal populations than those affected by a brain lesion. This distinction is essential when interpreting the resulting BOLD signal changes. While increases in BOLD activity are often taken to reflect compensatory engagement, they could alternatively indicate maladaptive or inefficient network recruitment. Future work using population tuning models or decoding approaches could help clarify the functional relevance of these activations and determine whether they truly reflect adaptive reorganization or instead a signature of disrupted processing.

Another intriguing aspect concerns the resting-state TMS-fMRI data (presented in the supplementary results). These results helped confirm the spatial specificity of the TMS effects, validating that stimulation targeted the intended network nodes even at rest. However, we observed that the direction of BOLD signal changes at rest contrasted with those seen during task performance. This apparent discrepancy has been reported in prior studies and may reflect state-dependent effects of TMS—that is, the neural response to TMS can differ markedly depending on whether the brain is at rest or engaged in a task. Another contributing factor could be the experimental design itself: we employed a block design during the resting-state acquisition and an event-related design during the task-based sessions. These methodological differences could influence the temporal integration of BOLD responses and the observed directionality of signal change (Chee *et al*., 2003; Tie *et al*., 2009). Together, these findings underscore the complexity of interpreting TMS-induced effects in fMRI data and highlight the need to consider both brain state and experimental context when drawing causal inferences.

## 6. Conclusion

Despite growing interest and significant advances in computational modelling, the causal mechanisms underlying neural network dynamics following a lesion remain incompletely understood. In this study, we employed concurrent TMS-fMRI to probe the neural mechanisms supporting functional reorganization across whole-brain networks, using a targeted perturbation approach.

Our findings offer neuroimaging evidence for the context-dependent nature of TMS effects, suggesting that such perturbations may preferentially engage specific neuronal populations depending on task demands. Moreover, we show that inducing focal, virtual lesions at different levels of the visual hierarchy leads to distinct patterns of dysregulation across visual networks.

Complementing classical lesion studies (Das *et al*., 2012), our results highlight the utility of TMS-fMRI coupling as a powerful tool for investigating ‘causal disconnectomics’—the causal relationships between localized disruptions and large-scale neural and behavioural consequences (Egger *et al*., 2021). This approach holds promise for precisely mapping how local perturbations propagate through brain networks, especially when assessed during active cognitive processing. Ultimately, such techniques may advance our understanding of brain resilience, compensation, and vulnerability in the face of focal damage.

## Acknowledgements

We thank Holly Bridge for her insightful comments on the manuscript. We would like to thank the MRI and neuromodulation facilities of the Human Neuroscience Platform of the Fondation Campus Biotech Geneva, for technical advice. This study was supported by the Bertarelli Foundation (Catalyst BC77O7 to FCH & ER), by the Swiss National Science Foundation (PRIMA PR00P3_179867 to ER), and by the Defitech Foundation (to FCH).

