## supplementary_materials for "Causal disconnectomics of motion perception: insights from TMS-induced BOLD responses"

**Supplementary online Materials**

**1.** **Motion processing localizer**: A standard MT localizer task was acquired during the first fMRI session and used to accurately and individually target the MT area. The screen displayed radially moving dots alternating with stationary dots (see e.g., (Sack et al., 2007)). A block design alternated six 15 s blocks of radial motion with six blocks featuring stationary white dots in a circular region on a black background. This region subtended 25° visual angle, with 0.5 dots per square degree. Each dot was 0.36° diameter. In the motion condition the dots repeatedly moved radially inward for 2.5 s and outward for 2.5 s, with 100% coherence, at 20°/s measured at 15° from the center. Participants were passively looking at the screen and were asked to focus on a fixation point located in the middle of the screen. The motion processing localizer was acquired using a GE-EPI sequence with 56 axial slices, slice thickness = 2.2 mm, in-plane resolution = 2.2 mm, TR = 2000 ms, TE = 30 ms, FOV = 242 mm, flip angle = 60°, GRAPPA = 2, Multiband Factor (MB) = 2.

**2. Additional MRI sequences for coregistration of TMS-fMRI data**

Functional and structural MR data, respectively acquired with the TMS-MRI coils and the 64-channel head coil, were found difficult to co-register due to different coil coverage, sensitivity and contrast. Therefore, two additional rapid high signal-to-noise, balanced, steady-state free precession (SSFP) sequences (Bieri and Scheffler, 2013) (96 axial slices, slice thickness = 2 mm, (gap 0.4 mm), in-plane resolution = 2 mm, TR = 5.24 ms, FOV = 256 mm, flip angle = 28°) were acquired with both the TMS-MRI coils and the body coil integrated into the scanner bore in order to resolve based on the same SSFP contrast the coil coverage mismatch within the registration processing pipeline.

**3. Supplementary results:** Relationship between awareness and accuracy

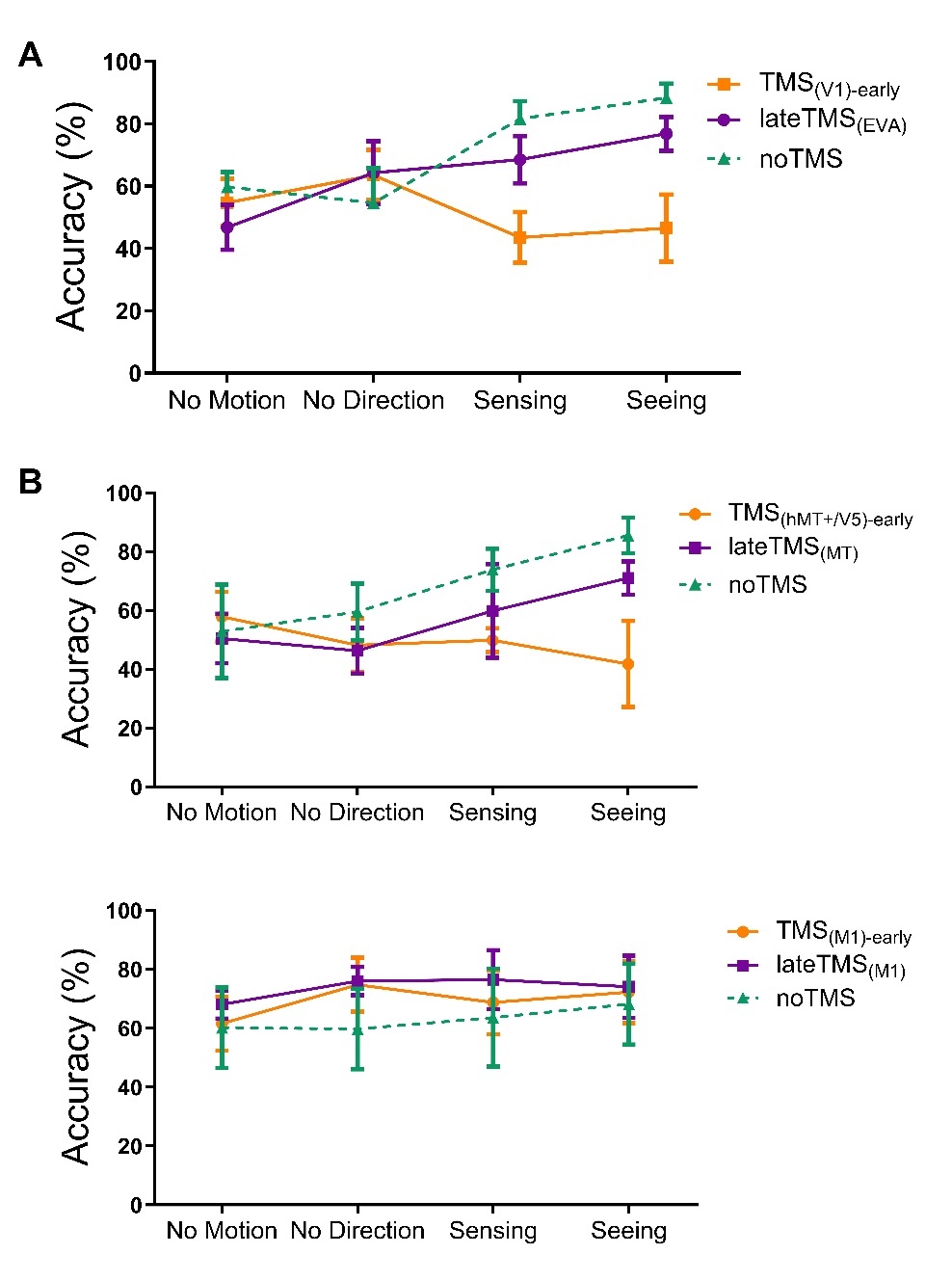

**Figure S1: *:*** *Relationship between behavioural accuracy and awareness of motion under noTMS (in green) TMS_early_ (in orange) and TMS_late_ (in purple) for TMS_(EVA)_ (A), TMS_(MT)_ (B) and TMS_(M1)_ (C).*

**4. Supplementary results: TMS induced activation at rest**

Figure S2 shows the conjunction map for TMS at rest induced by the two intensities. Statistically significant clusters (p < 0.001 cluster detection, p < 0.05 FWEc) are listed in Table 1 for both stimulated areas. For TMS_EVA_, significant activations were found in the right EVA, in the right medio-temporal cortex, the right insula and in the bilateral frontal cortex (Figure S2A and Table S1). To evaluated the effect of TMS intensities, the extracted parameters estimates (beta weights) were entered into a one-way ANOVA. It revealed a significant TMS condition effect F(2,30) = 11.38, p = 0.0002). Post hoc comparisons showed significant differences between TMS_High_ and noTMS (t(15) = 5.4, p < 0.001) and between TMS_Low_ and noTMS (t(15) = 2.86, p = 0.01), but no difference between TMS_High_ and TMS_Low_ (t(15) = 1.42, p = 0.17, Tukey’s multiple comparisons tests).

When TMS was delivered over the right MT, the conjunction analysis returned a significant activation cluster under the stimulated MT area. Additional remote clusters were found in the right EVA, the anterior and medial cingulate cortex as well as in the bilateral frontal eyed field (Figure S2B and Table S1). The local beta values extracted from the ipsilateral MT ROI demonstrated a significant TMS condition effect (F(2,30) = 10.36, p = 0.0004). As for the EVA stimulation, post hoc analyses showed significant differences between TMS_High_ and noTMS (t(15) = 3.87, p = 0.0015) and between TMS_Low_ and noTMS (t test: t(15) = 3.4, p = 0.004), but no difference between TMS_High_ and TMS_Low_ (t(15) = 0.2, p = 0.84).

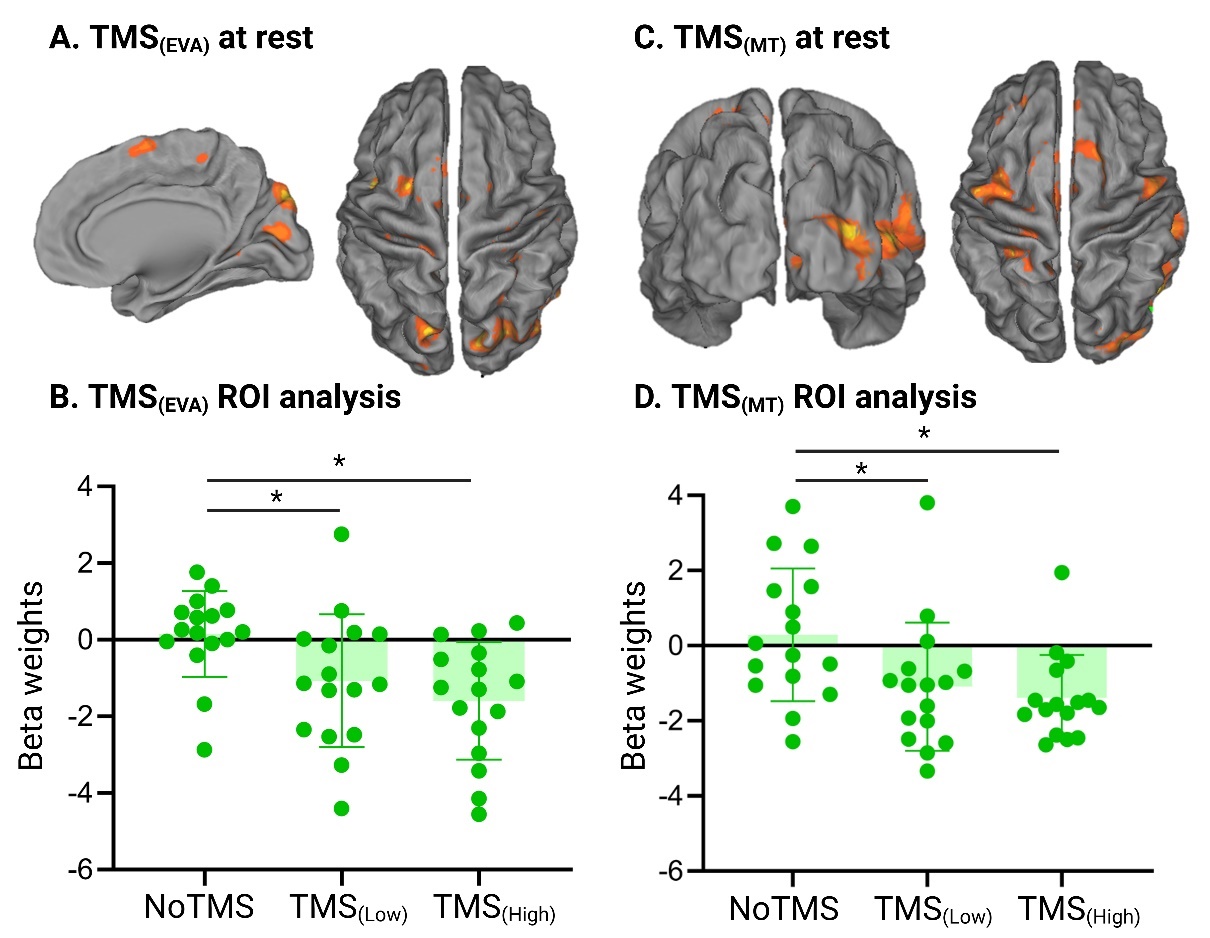

***Figure S2A:*** *Main effect of TMS_(EVA):_* ***2B****: Beta values for the three TMS conditions over EVA (one-way ANOVA);* ***2C****: Main effect of TMS_(MT)_;* ***2D:*** *Beta values for the three TMS conditions for MT (one-way ANOVA).*

**Supplementary Table S1: Significant clusters for TMS_(EVA)_ and TMS_(MT)_ at rest**

|  |  |  | MNI Coordinates | | |
| --- | --- | --- | --- | --- | --- |
| Region Label | **Extent** | **F-value** | **x** | **y** | **z** |
| TMS_(EVA)_ |  |  |  |  |  |
| R Lingual Gyrus (including EVA) | 2991 | 19.7683 | 20 | -64 | -2 |
| L + R Medial Frontal Gyrus | 80 | 13.0311 | 3 | 36 | 36 |
| L Middle Frontal gyrus | 96 | 11.435 | -30 | -2 | 52 |
| R MCC | 172 | 12.5029 | 10 | -44 | 56 |
| R + L Cuneus | 134 | 10.8385 | 18 | -74 | 36 |
| TMS_(MT)_ |  |  |  |  |  |
| R Middle Temporal Gyrus (including MT) | 5058 | 23.9567 | 24 | -48 | -2 |
| R Lingual Gyrus (including EVA) | 2782 | 18.32 | 4 | -91 | -4 |
| L + R Medial Frontal Gyrus | 217 | 16.7515 | -2 | 28 | 36 |
| L Middle Frontal Gyrus | 361 | 9.5591 | -38 | 0 | 64 |
| R Superior Frontal Gyrus | 416 | 15.7669 | 20 | 6 | 58 |
| L Posterior-Medial Frontal | 416 | 6.5147 | -6 | -6 | 58 |
| R & L IFG (p. Opercularis) | 50 | 12.2801 | 58 | 10 | 28 |
| R Rolandic Operculum | 50 | 6.0602 | 54 | 2 | 8 |

**5. Supplementary Table S2: Significant clusters for TMS_(EVA)_**

|  |  |  | MNI Coordinates | | |
| --- | --- | --- | --- | --- | --- |
| **Region Label** | **Extent** | **F-value** | **x** | **y** | **z** |
| *Visual stimulus effect* |  |  |  |  |  |
| R Medial Temporal Cortex | 157 | 17.9797 | 46 | -52 | 0 |
| R Superior Temporal Gyrus | 137 | 15.2246 | 44 | -42 | 18 |
| L Middle Frontal Gyrus | 227 | 16.5663 | -20 | -12 | 60 |
| R Middle Frontal Gyrus | 433 | 13.0977 | 24 | -5 | 58 |
| R Precuneus | 311 | 15.5128 | 10 | -60 | 52 |
| L Precuneus | 311 | 15.1918 | -14 | -46 | 64 |
| R Posterior-Medial Frontal | 433 | 15.121 | 8 | 12 | 54 |
| *TMS x Visual stimulus interaction* |  |  |  |  |  |
| L Calcarine Gyrus | 212 | 14.7565 | -10 | -84 | 14 |
| R Calcarine Gyrus | 250 | 14.4692 | 10 | -86 | 8 |
| R Anterior Cinculate Cortex | 226 | 14.1312 | 4 | 44 | 34 |

**6. Supplemetary Table S3: Significant clusters for TMS_(MT)_**

|  |  |  | MNI Coordinates | | |
| --- | --- | --- | --- | --- | --- |
| **Region Label** | **Extent** | **t-value** | **x** | **y** | **z** |
| *TMS effect* |  |  |  |  |  |
| R IFG (p. Triangularis) | 71 | 17.4877 | 32 | 14 | 32 |
| R Middle temporal Cortex | 400 | 16.419 | 44 | -71 | -9 |
| R Precuneus | 363 | 15.4239 | 20 | -76 | 46 |
| L Caudate Nucleus | 53 | 15.7919 | -14 | -6 | 24 |
| L ACC | 106 | 15.0247 | 0 | 22 | 36 |
| L Superior Medial Gyrus | 63 | 14.9261 | 2 | 48 | 32 |
| *TMS x Visual stimulus interaction* |  |  |  |  |  |
| R Linual Gyrus | 181 | 23.9332 | 22 | -90 | -6 |

**7. Supplementary Table S4: Local beta values (posthoc comparisons from the TMS * Visual stimulus interaction)**

|  |  | **T values** | **Corrected p values** |
| --- | --- | --- | --- |
| noTMS, Static | earlyTMS, Static | -0.25 | 1.00 |
|  | lateTMS, Static | -0.36 | 1.00 |
|  | noTMS, Moving | 1.2 | 1.00 |
|  | **earlyTMS, Moving** | **-3.2** | **0.04** |
|  | lateTMS, Moving | -1.8 | 0.7 |
| earlyTMS, Static | lateTMS, Static | -0.11 | 1.00 |
|  | noTMS, Moving | 1.2 | 1.00 |
|  | **earlyTMS, Moving** | **-3.6** | **0.02** |
|  | lateTMS, Moving | -1.5 | 1.00 |
| lateTMS, Static | noTMS, Moving | 1.3 | 1.00 |
|  | earlyTMS, Moving | -2.8 | 0.09 |
|  | lateTMS, Moving | -1.8 | 0.8 |
| noTMS, Moving | **earlyTMS, Moving** | **-4.2** | **0.003** |
|  | lateTMS, Moving | -2.8 | 0.09 |
| earlyTMS, Moving | lateTMS, Moving | 1.4 | 1.00 |

**8. Supplementary Table S5 (ICA EVA)**

| Component | Cluster region | Cluster extent | Peak MNI coordinate | | |
| --- | --- | --- | --- | --- | --- |
|  |  |  | x | y | z |
| **TMS-specific Networks** |  |  |  |  |  |
| IC1* | EVA (Bilat.) | 1843 | 4 | -85 | 20 |
| IC3 | EVA (Right hem.) | 1557 | 5 | -92 | 2 |
| IC7* | EVA (Bilat.) | 1243 | -1 | -81 | 18 |
| IC8 | EVA (Bilat.) | 1622 | -1 | -88 | 3 |
| **Other Networks** |  |  |  |  |  |
| IC4 | EVA (Left hem.) | 1363 | -8 | -89 | 16 |
| IC5 | Extrastriate (Bilat.) | 587 | 2 | -35 | 59 |
| IC6 | Med. Frontal (Bilat.) | 634 | -4 | 12 | 65 |

**9. Supplementary Table S6 (MT ICA)**

| **Component** | **Cluster region** | **Cluster extent** | **Peak MNI coordinate** | | |
| --- | --- | --- | --- | --- | --- |
|  |  |  | x | y | z |
| **TMS specific Networks** |  |  |  |  |  |
| IC1* | Medial Temporal Lobe (Right hem.) | 1343 | 48 | -70 | 12 |
|  | Inf. Temporal Lobe (Right hem.) | 187 | 60 | -68 | -6 |
|  | IPS (Right hem.) | 218 | 22 | -64 | 52 |
|  | Med. Frontal Lobe (Left hem.) | 138 | -4 | 58 | 58 |
|  | Sup. Parietal Lobe (Left hem.) | 127 | -6 | -64 | 60 |
| IC2 | Medial Temporal Lobe (Right hem.) | 348 | 43 | -73 | 16 |
|  | Sup. Parietal Lobe (Right hem.) | 377 | 6 | 59 | 56 |
| IC8* | Medial Temporal Lobe (Right hem.) | 687 | 39 | -63 | 15 |
| **Other Networks** |  |  |  |  |  |
| IC4 | EVA (Right hem.) | 420 | -3 | -78 | 2 |
| IC5 | IPS (Bilat.) | 722 | 31 | -68 | 55 |
|  | Sup. Frontal Gyrus (bilat.) | 683 | -13 | 69 | 12 |
| IC7 | Sup. Parietal Lobe (Bilat.) | 132 | -48 | -18 | 52 |
| IC6 | PreCuneus (Bilat.) | 456 | 4 | -64 | 58 |

**10. Supplementary Table S7** (paired t-tests for Sigma compared between TMS_(EVA)_ and TMS_(MT)_ networks)

| **Sparcity thresholds** | **t** | **df** | **p** |
| --- | --- | --- | --- |
| **0.05** | 0.084 | 14 | 0.23 |
| 0.1 | 0.121 | 14 | 0.24 |
| 0.15 | 0.31 | 14 | 0.45 |
| 0.2 | 0.52 | 14 | 0.26 |
| 0.25 | 0.84 | 14 | 0.16 |
| 0.3* | 2.01 | 14 | 0.03 |
| 0.35 | 1.89 | 14 | 0.06 |
| 0.4* | 2.01 | 14 | 0.03 |
| 0.45* | 2.02 | 14 | 0.04 |
| 0.5 | 1.45 | 14 | 0.17 |

**: Significant difference in smallworldness (Sigma) between EVA and MT networks.*

**11. Supplementary Table S8** (paired t-tests for Gamma compared between TMS_(EVA)_ and TMS_(MT)_ networks)

| **Sparcity thresholds** | **t** | **df** | **p** |
| --- | --- | --- | --- |
| 0.05 | 0.237 | 14 | 0.82 |
| 0.1 | 0.175 | 14 | 0.86 |
| 0.15 | 0.128 | 14 | 0.9 |
| 0.2 | 0.26 | 14 | 0.8 |
| 0.25 | 0.5 | 14 | 0.62 |
| 0.3 | 0.713 | 14 | 0.5 |
| 0.35 | 0.845 | 14 | 0.42 |
| 0.4 | 0.946 | 14 | 0.36 |
| 0.45 | 1.089 | 14 | 0.2 |
| 0.5 | 1.089 | 14 | 0.28 |

**12. Supplementary Table S9** (paired t-tests for Lambda compared between TMS_(EVA)_ and TMS_(MT)_ networks)

| **Sparcity thresholds** | **t** | **df** | **p** |
| --- | --- | --- | --- |
| 0.05 | 0.452 | 14 | 0.66 |
| 0.1 | -0.72 | 14 | 0.49 |
| 0.15(*) | -1.9 | 14 | 0.08 |
| 0.2* | -2.03 | 14 | 0.05 |
| 0.25* | -2.16 | 14 | 0.04 |
| 0.3 | -1.45 | 14 | 0.17 |
| 0.35 | -1.2 | 14 | 0.25 |
| 0.4 | -0.75 | 14 | 0.47 |
| 0.45 | -0.8 | 14 | 0.43 |
| 0.5 | -0.77 | 14 | 0.46 |

**: Significant difference in Characteristic Path Length (Lambda) between EVA and MT networks.*

**13. Supplementary Table S10: Multiple linear regressions**

| **Models** | Analysis of Variance | SS | **DF** | **MS** | **F(DFn,DFd)** | **P value** |
| --- | --- | --- | --- | --- | --- | --- |
| **TMS_(EVA)_ early** | Regression | 2834 | 6 | 472.4 | F (6, 7) = 1.007 | P=0.4886 |
|  | BetaV1 | 144.4 | 1 | 144.4 | F (1, 7) = 0.3077 | P=0.5964 |
|  | IC1 | 65.44 | 1 | 65.44 | F (1, 7) = 0.1394 | P=0.7199 |
|  | IC7 | 1999 | 1 | 1999 | F (1, 7) = 4.260 | P=0.0779 |
|  | meanLambda | 406.9 | 1 | 406.9 | F (1, 7) = 0.8670 | P=0.3828 |
|  | meanGamma | 58.02 | 1 | 58.02 | F (1, 7) = 0.1236 | P=0.7355 |
|  | MeanSigma | 2.066 | 1 | 2.066 | F (1, 7) = 0.004402 | P=0.9490 |
|  | Residual | 3285 | 7 | 469.3 |  |  |
|  | Total | 6119 | 13 |  |  |  |
| **TMS_(MT)_ early** | Regression | 3287 | 6 | 547.9 | F (6, 6) = 1.719 | P=0.2634 |
|  | BetaMT | 2.931 | 1 | 2.931 | F (1, 6) = 0.009196 | P=0.9267 |
|  | IC1 | 350.6 | 1 | 350.6 | F (1, 6) = 1.100 | P=0.3347 |
|  | IC8 | 25.8 | 1 | 25.8 | F (1, 6) = 0.08095 | P=0.7856 |
|  | meanLambda | 0.08906 | 1 | 0.08906 | F (1, 6) = 0.0002794 | P=0.9872 |
|  | **meanGamma** | **2115** | **1** | **2115** | **F (1, 6) = 6.636** | **P=0.0420** |
|  | MeanSigma | 18.98 | 1 | 18.98 | F (1, 6) = 0.05956 | P=0.8153 |
|  | Residual | 1913 | 6 | 318.8 |  |  |
|  | Total | 5200 | 12 |  |  |  |
| **TMS_(EVA)_ late** | Regression | 381.9 | 6 | 63.65 | F (6, 4) = 5.470 | P=0.0610 |
|  | BetaEVA | 4.257 | 1 | 4.257 | F (1, 4) = 0.3659 | P=0.5779 |
|  | IC7 | 48.24 | 1 | 48.24 | F (1, 4) = 4.146 | P=0.1114 |
|  | IC3 | 43.62 | 1 | 43.62 | F (1, 4) = 3.749 | P=0.1249 |
|  | meanLambda | 33.91 | 1 | 33.91 | F (1, 4) = 2.915 | P=0.1630 |
|  | meanGamma | 0.8721 | 1 | 0.8721 | F (1, 4) = 0.07495 | P=0.7978 |
|  | MeanSigma | 0.6260 | 1 | 0.6260 | F (1, 4) = 0.05380 | P=0.8280 |
|  | Residual | 46.54 | 4 | 11.64 |  |  |
|  | Total | 428.4 | 10 |  |  |  |
| **TMS_(MT)_ late** | Regression | 334.7 | 6 | 55.78 | F (6, 6) = 0.9866 | P=0.5063 |
|  | BetaMT | 7.344 | 1 | 7.344 | F (1, 6) = 0.1299 | P=0.7309 |
|  | IC1 | 296.9 | 1 | 296.9 | F (1, 6) = 5.250 | P=0.0618 |
|  | IC8 | 132.4 | 1 | 132.4 | F (1, 6) = 2.342 | P=0.1768 |
|  | meanLambda | 88.09 | 1 | 88.09 | F (1, 6) = 1.558 | P=0.2585 |
|  | meanGamma | 51.87 | 1 | 51.87 | F (1, 6) = 0.9174 | P=0.3751 |
|  | MeanSigma | 16.00 | 1 | 16.00 | F (1, 6) = 0.2830 | P=0.6139 |
|  | Residual | 339.2 | 6 | 56.54 |  |  |
|  | Total | 673.9 | 12 |  |  |  |
